# Transplacental protection against asthma by maternal treatment with a bacterial-derived immunomodulatory agent

**DOI:** 10.1101/232231

**Authors:** Kyle T. Mincham, Naomi M. Scott, Jean-Francois Lauzon-Joset, Jonatan Leffler, Alexander N. Larcombe, Philip A. Stumbles, Sarah A. Robertson, Christian Pasquali, Patrick G. Holt, Deborah H. Strickland

**Affiliations:** Telethon Kids Institute, University of Western Australia, Subiaco, Western Australia, Australia.; Health, Safety and Environment, School of Public Health, Curtin University, Perth, Western Australia, Australia.; School of Veterinary and Life Sciences, Murdoch University, Perth, Western Australia, Australia.; School of Paediatrics and Child Health, University of Western Australia, Subiaco, Western Australia, Australia.; Robinson Research Institute and School of Medicine, University of Adelaide, Adelaide, South Australia, Australia.; OM Pharma, SA Geneva, Geneva, Switzerland.

## Abstract

Studies in European and US farming populations have documented major reductions in asthma prevalence in offspring of mothers exposed to microbial breakdown products present in farm dusts and unprocessed foods. This was associated with enhancement of innate immune competence in the offspring. We sought to (i) identify a safe therapeutic that would reproduce these immunomodulatory effects in a murine model, (ii) elucidate underlying mechanism(s)-of-action, and (iii) develop a scientific rationale for progressing this approach to human trials. We demonstrate in mice that maternal treatment during pregnancy with the microbial-derived immunomodulator OM85, which has been used clinically in adults and children in Europe for >30 years for bolstering resistance to infection-associated airways inflammation, markedly reduces the susceptibility of the offspring of treated mothers to development of experimental atopic asthma. We identify bone marrow precursors of the dendritic cell populations responsible for airway mucosal immune surveillance as the primary targets for the asthma-protective effects of maternal OM85 treatment in the offspring.

## INTRODUCTION

A series of prospective birth cohort studies on the children of European traditional farming families^1, 2^, now replicated in US studies contrasting Amish and Hutterite farming populations^3^, have identified striking asthma-protective properties of oral and inhalation exposure to benign microbial stimuli present in dusts from farm barns. The target for these exposures in the offspring appears to be the innate immune system, and involves modulation of both immunoregulatory and effector cell function (s)^3-6^ resulting in markedly reduced susceptibility to the asthma-promoting effects of common respiratory allergies. The temporal window during which these environmental stimuli exert their immunomodulatory effects spans the period when the developing immune system is undergoing postnatal functional maturation, but susceptibility to these effects also appears particularly high during prenatal development as demonstrated by the strong impact of maternal microbial exposures during pregnancy on ensuing asthma resistance in their offspring ^1,7^.

We posited that if these benign environmental exposure effects could be reproduced by a therapeutic that could be safely administered during pregnancy, then this could open up novel possibilities for primary prevention of asthma. With this in mind we have recently completed a proof-of-concept study in pregnant mice with a microbial-derived therapeutic product OM85, which has been in widespread use in Europe in infants and adults for >30 years for boosting resistance to airways inflammation and attendant wheezing symptoms associated with lower respiratory infections^8-12^. In preliminary investigations to establish the safety of OM85 use during pregnancy we demonstrated that maternal treatment with this agent enhanced homeostatic control of innate immune and inflammatory functions in gestational tissues at baseline and in the face of challenge with microbial pathogens including live influenza infection and the bacterial mimic LPS. Specifically, OM85 treatment attenuated inflammatory symptoms (which are typically exaggerated during pregnancy) and protected against fetal growth restriction and/or pregnancy termination which can follow maternal infection^13^. In the study presented here we focus on the effects of maternal OM85 treatment during healthy pregnancy on the immunocompetence of offspring during the postnatal weanling period when immune functions are typically developmentally compromised. In this regard, we focus on the effects of maternal OM85 treatment on the capacity of offspring to regulate airways inflammatory responses associated with development of experimental atopic asthma during the weanling period, which in humans represents the age range at highest risk for initiation of what can be life-long asthma^14^.

## RESULTS

### Experimental model of atopic asthma in sensitized weanling mice: study rationale

For this study we utilized an experimental system developed for induction of Th2-associated cell-mediated inflammation in the conducting airway mucosa in adult rodents, as a model for the main lesional site in human asthma. Additional (albeit less extensive) inflammation also develops in peripheral lung tissue, but the relative contribution of this to airflow limitation in the asthmatic state is uncertain. The principal features of this model, focusing mainly on the airway mucosa, are illustrated in Fig. S1A. Aeroallergen delivered to the airways of presensitized animals via large-droplet aerosol is captured by resident mucosal dendritic cells (DC) that are functionally quiescent in the steady-state (as marked by low-modest IAIE expression) and are specialised for antigen sampling only, which they subsequently transport to airway draining lymph nodes (ADLN) for presentation to allergen-specific T-memory cells^15-17^. The resultant T-cell response generates a mixture of T-effector-memory (T^m^eff) and T-regulatory (Treg) cells in proportions determined via DC programming. Representatives of these populations traffic back to the airway mucosa, where they encounter resident mucosal DC which have recently acquired aeroallergen, and bidirectional interactions between these three cell populations *in situ* determine the intensity and duration of the ensuing T-cell dependent inflammatory response within the airway mucosa^18, 19^ In particular, the capacity for local activation of incoming T^m^effs is limited via the suppressive effects of Tregs on surface IAIE and CD86 expression by mucosal conventional DC (cDC)^15, 19-21^.

The principal DC population involved comprises the network of cDC within the airway mucosa, that are responsible for the major aspects of local immune surveillance^15^. Plasmacytoid DC (pDC) have also been implicated in this process^22^, particularly in relation to pathogen surveillance^23^. The airway mucosal cDC population has the property of uniquely rapid turnover in the steady-state with 85% of the resident population turning over every ~24 hours, being continuously depleted by migration of antigen-bearing cells to ADLN and simultaneously replenished via incoming precursors recruited from bone marrow^24^. This orderly and highly dynamic process is rapidly accelerated during airway challenge events, during which cDC numbers can expand markedly within the airway epithelium and ADLN^15,25-27^. This airway mucosal cDC network is developmentally compromised in immature humans^28, 29^ and experimental animals^30, 31^ and this partially explains the high risk of respiratory infections and aeroallergen sensitization associated with the infant period^32^. Our hypothesis underlying this murine study is that maternal OM85-treatment during pregnancy can enhance the functional maturation of this mucosal immune surveillance system in their offspring, and as a result reduce susceptibility to initiation of inflammatory airway disease during the high-risk early postnatal period.

To test this hypothesis, we have utilised a variant of the above mentioned asthma model, modified from earlier studies assessing farm-related exposures^33^, involving sensitization of 21 day old weanling BALB/c mice to ovalbumin (OVA) employing a prime-boost schedule followed by subsequent airways challenge with aerosolised OVA (Fig. S1B, C). Details of the ensuing response are discussed below.

### Aeroallergen-induced cellular response in the airways: baseline characteristics

Sensitized animals display high levels of OVA-specific serum IgE 24 hours following repeated OVA-aerosol challenge (Fig. 1A). The challenged animals display gross hypertrophy of ADLN involving in particular T-cells (see below), which is accompanied by intense inflammatory cell infiltration into the airways encompassing eosinophils, neutrophils and lymphocytes detectable by BAL (Fig. 1B), increased levels of Th2 cytokines in lung homogenates (not shown), and airways hyperresponsiveness (AHR) manifesting as increased airways resistance (R_aw_) to methacholine (MCh; Fig. 1C).

**Fig. 1.**
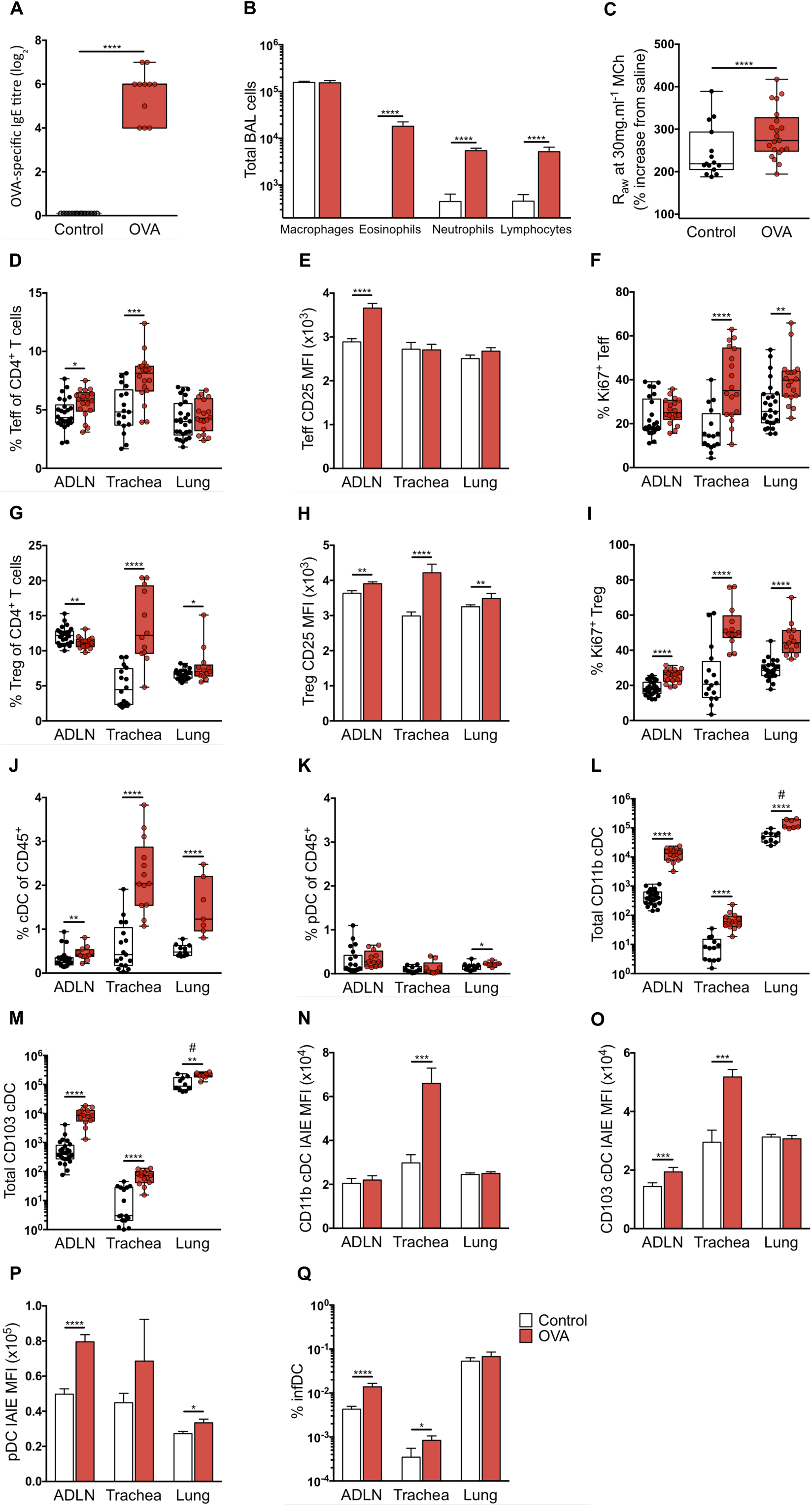
Baseline airways cellular characteristics following early life sensitisation and challenge. (**A**) Serum titres of OVA-specific IgE as measured by *in vivo* passive cutaneous anaphylaxis assay. (**B**) Absolute numbers of macrophages, eosinophils, neutrophils and lymphocytes as determined by bronchoalveolar lavage (BAL) 24 hours post challenge. (**C**) Airways hyperresponsiveness to methacholine (MCh) challenge at a dose of 30mg.ml^-1^ MCh. CD3^+^CD4^+^CD25^+^FoxP3^-^ T-effector cell (T^m^eff) proportion (**D**), CD25 mean fluorescence intensity (MFI) expression (**E**) and proliferation (**F**) within parathymic and mediastinal (airways draining) lymph nodes (ADLN), trachea and peripheral lung samples. CD3^+^CD4^+^CD25^+^FoxP3^+^ T-regulatory cell (Treg) proportion (**G**), CD25 mean fluorescence intensity expression (**H**) and proliferation (**I**) within ADLN, trachea and peripheral lung samples. IAIE+F4/80^-^CD11c^+^ conventional DCs (cDCs) (**J**) and IAIE^+^Ly6G/C^lo^F4/80^-^ CD11c^+^CD11b^-^B220^+^ plasmacytoid DCs (pDCs) (**K**) as a proportion of total CD45^+^ leukocytes within ADLN, trachea and peripheral lung samples. Absolute numbers of CD11b^+^ (**L**) and CD103^+^ (**M**) cDCs within ADLN, trachea and peripheral lung samples. MFI of IAIE expressed by CD11b^+^ cDCs (**N**), CD103^+^ cDCs (**O**) and pDCs (**P**) within ADLN, trachea and peripheral lung samples. (**Q**) Inflammatory DCs (infDC) as a proportion of total bone marrow. All data presented as OVA sensitised and aerosol challenged (red) offspring versus naïve controls (white) and displayed as box and whisker plot showing min to max or bar graph showing mean ± SEM of n≥4 independent experiments. Total peripheral lung cells displayed as cells per milligram of tissue (#). Data points represent individual animals. Statistical significance was determined using Student’s t-test or Mann Whitney test (A-B, E-Q) or two-way ANOVA followed by Sidak’s multiple comparison test (C) and presented as *p<0.05, **p<0.01, ***p<0.001, ****p<0.0001.

Further characterization of the phenotype of the cellular response within the airways compartment by multi-colour flow cytometry (see Methods for gating strategies) revealed significant increases in the total cellularity of parathymic and mediastinal ADLN and trachea, with no observable difference in peripheral lung (Fig. S1D). This cellular response was dominated by changes in the CD3^+^ T-cells (Fig. S1E) and especially the CD4^+^ T-cell compartment (Fig. S1F). These changes in particular involved increases in the numbers (Fig. S1G), proportions (Fig. 1D) and activation status (Fig. 1E, F) of CD3^+^CD4^+^CD25^+^FoxP3^-^T^m^eff cells within the ADLNs and in tracheal tissue, with much smaller parallel changes in peripheral lung parenchyma (Fig. 1D) which is consistent with deposition of the bulk of aerosol droplets in the large and central airways. ADLN T^m^effs displayed a heightened state of activation (Fig. 1E), whereas lung and especially tracheal T^m^effs demonstrated high levels of Ki67 expression suggesting very recent (possibly local) proliferation (Fig. 1F; see corresponding cDC data below). In conjunction with the CD25+FoxP3^-^ T^m^eff response, a decrease in CD3^+^CD4^+^CD25^+^FoxP3^+^ Tregs within the T-cell compartment of ADLNs was observed, accompanied by a large increase within trachea and a smaller (but significant) increase in peripheral lung tissues (Fig. 1G), presumably derived by migration of these cells from ADLN. Characterisation of Treg function-associated markers revealed increased CTLA-4^+^ Tregs within trachea (74.47 ± 3.78; 46.41 ± 6.05) and peripheral lung (51.16 ± 2.50; 34.9 ± 2.54) compared to naïve controls (Fig. S1H). Furthermore, an increase in CD69^+^ Tregs was identified within peripheral lung samples (15.68 ± 0.53; 12.98 ± 1.54), while CD69^+^ Tregs were reduced within the ADLNs (28.19 ± 0.69; 32.5 ± 1.19) compared to naïve controls (Fig. S1I). Airways Tregs additionally displayed enhanced CD25 expression (Fig. 1H) and proliferative capacity (Fig. 1I) following OVA challenge compared to naïve controls.

Forerunner studies from our lab^15, 19, 34^ suggest that in the early stage of recall responses to inhaled antigen, the limiting factor determining the efficiency of generation of airway mucosal homing Tregs is the efficiency of DC-mediated transport of antigen-specific signals from the airway to ADLN. We therefore turned our attention to characterising the myeloid cell populations localised within the airways of early life OVA sensitised and aerosol challenged animals. For these analyses, IAIE^+^F4/80^-^CD11c^+^ conventional DCs (cDC) were subdivided into CD11b^+^CD103^-^ and CD11b^-^CD103^+^ populations, representing the two dominant cDC subsets localised within the airways, having specialised roles in immunogenic and tolerogenic responses respectively^35^. We additionally quantified IAIE^+^Ly6G/C^lo^F4/80^-^ CD11c^+^B220^+^CD11b^-^ pDC which represented a much smaller proportion of the CD45^+^ population compared to cDC (Fig. 1J, K). Consecutive aerosol challenges of pre-sensitised mice induces a minor response in pDC in peripheral lung only (Fig. 1K), in contrast to a significant influx of both cDC subsets across all airways tissues sampled, in particular the tracheal mucosa and its ADLN where numbers of these cells displayed a logfold increase (Fig. 1L, M). Characterisation of cDC based on expression levels of surface IAIE demonstrated marked enhancement in expression intensity across the entire cDC population within the trachea following challenge (Fig. 1N, O). This observation is consistent with allergen-driven functional maturation *in situ* of these cells from strict antigen-sampling to antigen-presentation phenotype as previously observed^15, 16^, and this provides a plausible mechanism for local activation of T^m^effs within the mucosa during repeated challenge (Fig. 1D, F). In parallel, ADLN displayed modest IAIE upregulation in the CD103^+^ subset, while both subsets remained at baseline in the peripheral lung (Fig. 1N, O). Furthermore, upregulation of pDC IAIE expression occurred both in ADLN and peripheral lung (Fig. 1P). Additionally, we observed increased numbers post challenge of rare IAIE^+^F4/80^int^Ly6G/C^hi^CD11b^+^CD11c^+^ inflammatory DCs (infDC) which have been implicated in driving Th2-mediated inflammation to antigen exposure^36, 37^, within ADLN and tracheal tissue (Fig. 1Q).

### Role of the bone marrow in the experimental asthma response

The granulocytic and DC subsets identified above as participants in the airways response to aerosol challenge in sensitized mice are derived from bone marrow, and we posited that (as inferred from earlier studies on eosinophils in aeroallergen challenged human asthmatics^38^) the dynamic changes detailed in these populations in our murine asthma model above should be mirrored by changes in respective bone marrow precursor populations.

The scheme in Fig. S2 summarizes current understanding of the interrelationships between relevant bone marrow progenitor compartments in the development pathway leading to production of granulocytes, pDC and cDC. We assessed the impact of repeated challenge with aeroallergen on the size of relevant compartments at or beyond the myeloid precursor (MP) stage, employing multicolour flow cytometry, targeting the markers shown. These analyses (Fig. 2A-D) demonstrate firstly that early myeloid precursor compartments up to and including the granulocyte-macrophage progenitor (GMP) population, which are a major source of both DC and granulocyte populations^39, 40^, and the macrophage-dendritic cell progenitor (MDP) compartment which is committed to pDC and cDC production^41-43^, expand significantly in response to repeated aeroallergen challenge of sensitized animals. This finding is consistent with the data shown in Fig. 1, demonstrating the buildup of these populations in the challenged airways. Beyond this stage the cDC compartment appeared reduced relative to baseline (Fig. 2F), and this may be expected in light of the accumulation of these cells in tracheal mucosa and at their ultimate destination in ALDN, which displays a logfold increase in cDC numbers post-challenge (Fig. 1L, M).

**Fig. 2.**
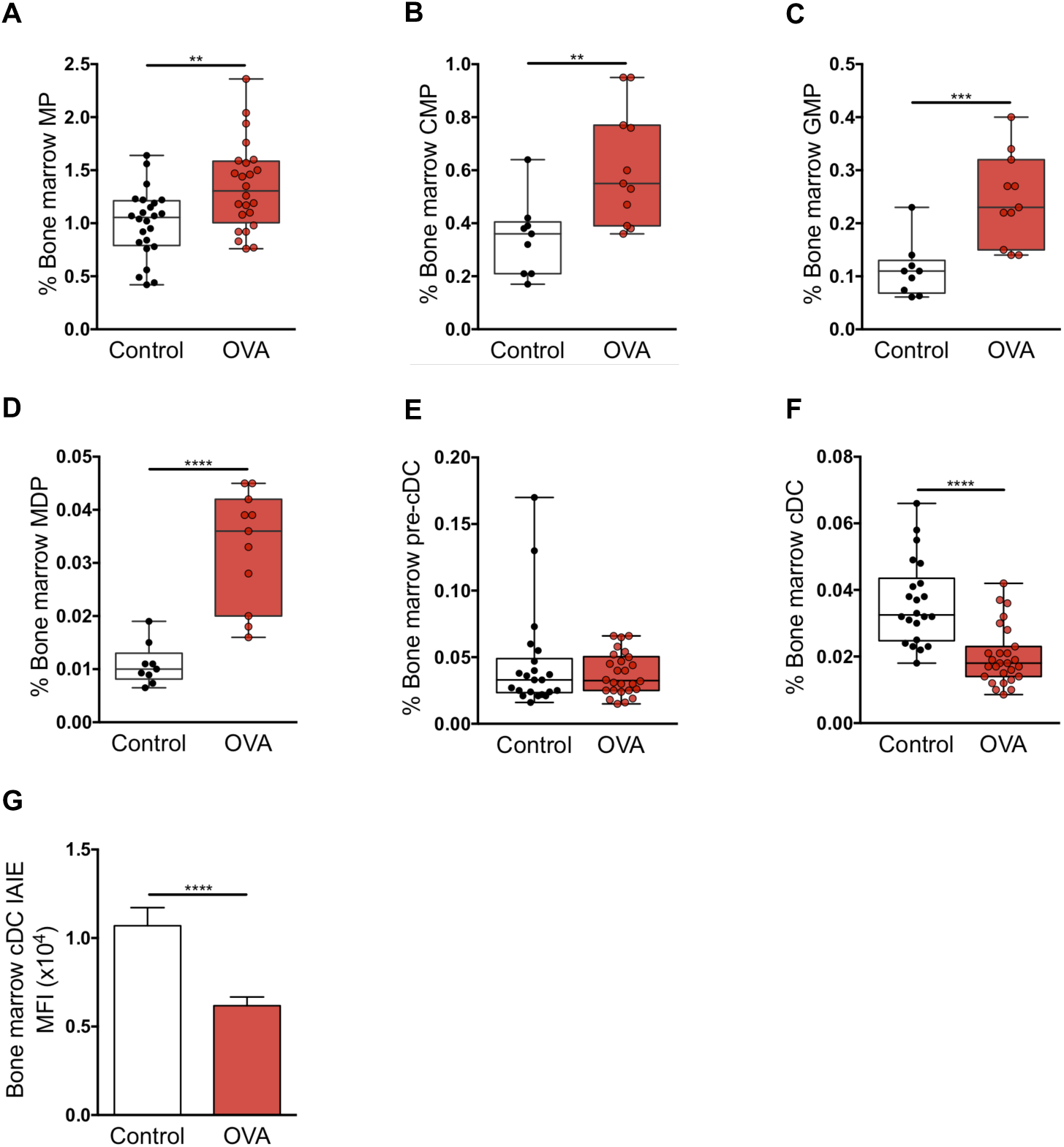
Baseline bone marrow response following early life sensitisation and challenge. Bone marrow Lin^-^IL7-Rα^-^c-Kit+Sca-1^-^ myeloid progenitors (MP) (**A**), Lin^-^IL7-Rα^-^c-Kit^+^Sca-1^-^CD16/32^lo/-^CD34^lo/-^ common myeloid progenitors (CMP) (**B**), Lin^-^IL7-Rα^-^c-Kit^+^Sca-1^-^ CD16/32^hi^CD34^+^ granulocyte-macrophage progenitors (GMP) (**C**), Lin^-^IL7-Rα^-^c-Kit^+^Sca-1^-^ CD16/32^hi^CD34^+^ CX_3_CR1^+^Flt-3^+^ macrophage-dendritic cell progenitors (MDP) (**D**), CD11c^+^CD11b^+^IAIE^-^ pre-cDCs (**E**), CD11c^+^CD11b^+^IAIE^+^ cDCs (**F**) and MFI of IAIE expression on cDCs (**G**) in OVA sensitised and aerosol challenged (red) offspring versus naïve controls (white). Cellular data presented as proportion of bone marrow and findings are identical if populations are expressed as total bone marrow cell numbers. Data displayed as box and whisker plot showing min to max or bar graph showing mean ± SEM of n≥8 independent experiments. Data points represent individual animals. Statistical significance was determined using Student’s t-test or Mann Whitney test and presented as **p<0.01, ***p<0.001, ****p<0.0001.

Furthermore, cDCs remaining within the bone marrow post challenge displayed reduced IAIE surface expression relative to baseline controls (Fig. 2G), which may indicate preferential recruitment of cDC from the more functionally mature end of the developmental spectrum.

### Maternal OM85 treatment during pregnancy: effects on experimental asthma susceptibility in sensitized offspring

We posited that treatment of pregnant mice with the microbial-derived immunomodulatory agent OM85 would enhance the resistance of their offspring to development of experimental atopic asthma during the early post-weaning period. To test this hypothesis, we utilised an OM85 treatment protocol we have recently demonstrated to protect pregnant mice and their fetuses against the toxic effects of bacterial and viral infections^13^, comprising oral administration of OM85 from gestation day 9.5-17.5, followed by natural delivery of offspring 2-3 days later. Age-matched offspring from OM85 treated and untreated control mothers were sensitized at weaning (21 days of age) and aerosol challenged as per Fig. S1B and C, and their airways responses compared (Fig. 3). As previously demonstrated, early life OVA sensitisation and ensuing aerosol challenge initiates granulocytic and lymphocytic infiltration of the airways, and these cellular responses (Fig. 3A), together with accompanying development of AHR to MCh (Fig. 3B), were markedly attenuated in the offspring of mice treated during pregnancy. Of note, treatment did not affect OVA-specific IgE levels (log2 titres 5.44 ± 0.21 and 5.54 ± 0.34 in treated versus untreated groups respectively), implying that OM85 treatment influences mechanism(s) downstream of sensitization *per se*.

**Fig. 3.**
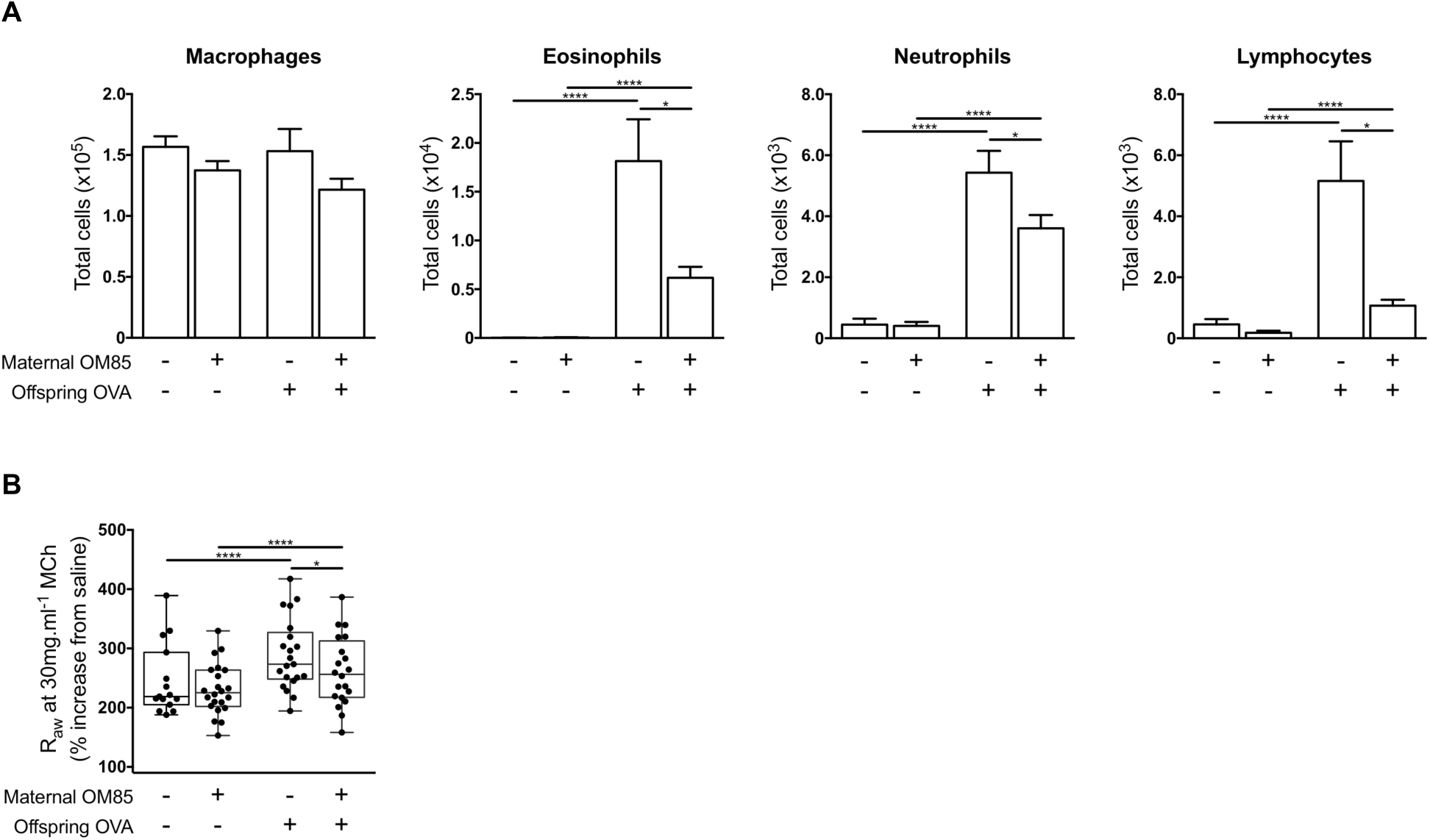
Attenuation of OVA-induced clinical disease following maternal OM85 pretreatment. (**A**) Absolute numbers of macrophages, eosinophils, neutrophils and lymphocytes within BAL 24 hours post challenge. (**B**) AHR to 30mg.ml^-1^ MCh challenge. Data presented for OVA sensitised and aerosol challenged offspring versus naïve controls from OM85 treated/untreated mothers and displayed as box and whisker plot showing min to max or bar graph showing mean ± SEM of n≥4 independent experiments. Data points represent individual animals. Statistical significance was determined using Student’s t-test (A) or two-way ANOVA followed by Sidak’s multiple comparison test (B) and presented as *p<0.05, ****p<0.0001.

### Treg function in offspring as a potential target for maternal OM85 treatment effects

Previous studies from our lab and others have identified mucosal homing Tregs as a potential target for OM85-mediated treatment effects in adult non-pregnant^44, 45^ and pregnant^13^ animals, prompting an initial focus on this population. We accordingly phenotyped the T-cell response within the airways compartment using multi-colour flow cytometry. The magnitude of the overall CD4^+^ T-cell response to OVA aerosol was reduced in the treatment group, particularly in the tracheal mucosa (Fig. 4A, B). Following challenge, the proportion of Tregs within the ADLN CD4+ T-cell population declined in both groups (Fig. 4C) and correspondingly increased in respective tracheal tissues (Fig. 4F), consistent with their migration to the airway mucosal challenge site. However, both the proportional decline in Tregs in ADLN and the corresponding increase in trachea were significantly higher in the OM85-treatment group (Fig. 4F). Treg: T^m^eff ratios post challenge remained higher in ADLN in offspring from treated mothers (Fig. 4E) but did not differ significantly between the groups in trachea (data not shown). Furthermore, the relative expression levels of Treg function-associated molecules CD25 (Fig. 4G), CTLA-4 (Fig. 4I) and FoxP3 (Fig. 4K), along with activation/proliferation-associated markers CD69 (Fig. 4J) and Ki67 (Fig. 4H), were significantly elevated in tracheal Tregs from the treated group, consistent with activation and enhanced functionality. Similar patterns were also observed for Treg populations from peripheral lung tissues (Fig. S3).

**Fig. 4.**
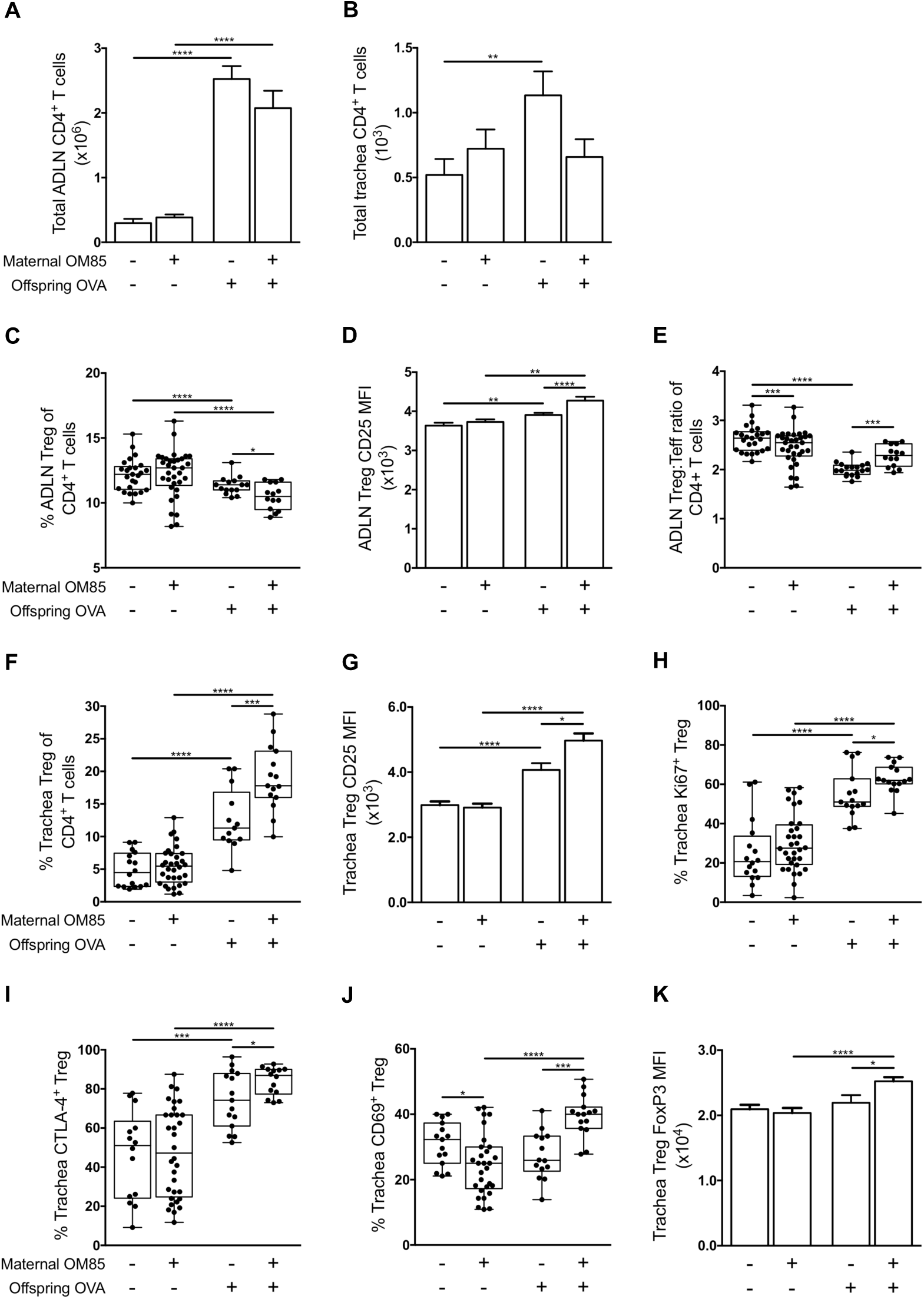
Maternal OM85 treatment promotes upregulation of Treg suppressive phenotype. Absolute numbers of CD3^+^CD4^+^CD8^-^ T-cells in ADLN (**A**) and trachea (**B**) samples. (**C**) CD3^+^CD4^+^CD25^+^FoxP3^+^ Tregs as a proportion of the CD4^+^ T-cell pool within ADLNs. (**D**) MFI of Treg CD25 expression in ADLN samples. (**E**) Treg-to-Teff ratio within the CD4^+^ T-cell pool of ADLNs. Treg proportion within the CD4^+^ T-cell pool (**F**), MFI of CD25 (**G**) and proliferation (**H**) within the trachea. Proportion of Tregs expressing intracellular CTLA-4 (**I**) and extracellular CD69 (**J**) within trachea. (**K**) MFI of intracellular FoxP3 expression in trachea Tregs. Data presented for OVA sensitised and aerosol challenged offspring versus naïve controls from OM85 treated/untreated mothers and displayed as box and whisker plot showing min to max or bar graph showing mean ± SEM of n≥9 independent experiments. Data points represent individual animals. Statistical significance was determined using Student’s t-test or Mann Whitney test and presented as *p<0.05, **p<0.01, ***p<0.001, ****p<0.0001.

### Maternal OM85 pre-treatment modulates the functional phenotype of airway-associated DC populations in offspring

The accumulation of cDC in airway-associated tissue compartments in response to airways challenge was generally reduced in the treated group, and these differences were statistically significant for CD103^+^ cDC in the trachea, and for CD11b^+^ cDC in ADLN and lung (Fig. 5A, B). However, the most notable finding related to cDC maturational status as measured by surface IAIE expression, which was reduced at baseline in both CD103^+^ and CD11b^+^ subsets in ADLN (Fig. 5C). Moreover, the antigen-induced surge in IAIE expression levels on both subsets in the rapidly turning over cDC population in the tracheal mucosa, which is a hallmark of functional activation of these cells, was likewise attenuated (Fig. 5D). This contrasted with the picture in the peripheral lung (Fig. 5E), which is dominated by cDC populations with much longer half-lives, and which displayed minimal upregulation of IAIE in response to challenge. However as noted above, aerosol challenge does elicit a small but significant increase in pDC in peripheral lung tissue, and this response was attenuated in the treated group (Fig. 5F, G). We also screened the groups for treatment effects on the rare inflammatory DC subset in airway tissues, however none were detected (data not shown).

**Fig. 5.**
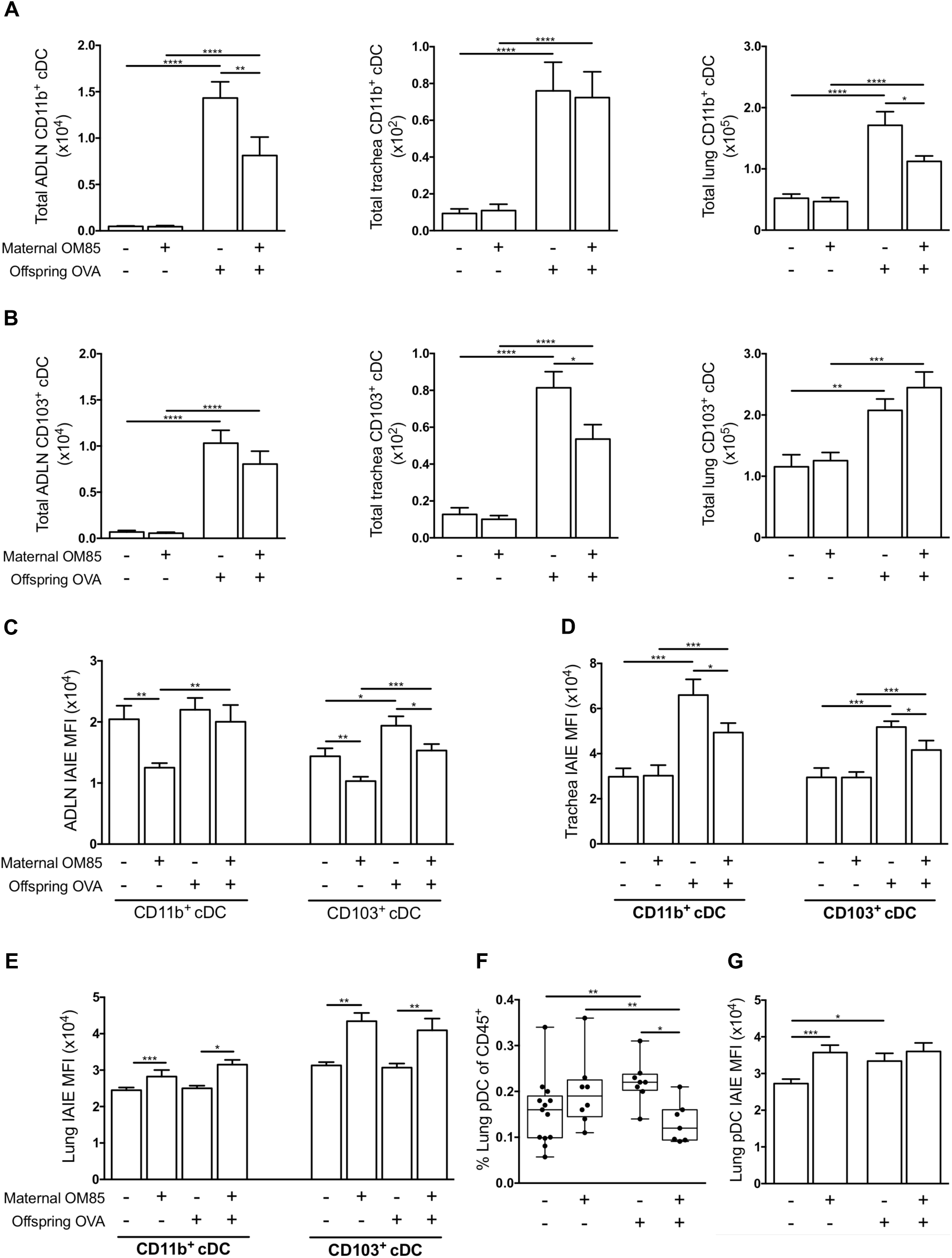
Diminished airways mucosal DC response following maternal OM85 pretreatment. Absolute numbers of CD11b^+^ (**A**) and CD103^+^ (**B**) cDCs within ADLN, trachea and peripheral lung. MFI of IAIE expression on CD11b^+^ and CD103^+^ cDCs within ADLN (**C**), trachea (**D**) and peripheral lung (E). Peripheral lung pDC proportion (**F**) and IAIE MFI (**G**). Data presented for OVA sensitised and aerosol challenged offspring versus naïve controls from OM85 treated/untreated mothers and displayed as box and whisker plot showing min to max or bar graph showing mean ± SEM of n≥4 independent experiments. Total peripheral lung cells displayed as cells per milligram of tissue. Data points represent individual animals. Statistical significance was determined using Student’s t-test or Mann Whitney test and presented as *p<0.05, **p<0.01, ***p<0.001, ****p<0.0001.

### Offspring bone marrow as the primary target for maternal OM85 treatment effects

The final series of experiments tested the hypothesis that maternal OM85 treatment mediated effects on cellular immune function(s) in offspring respiratory tract tissues in this model may be associated with upstream effects on relevant precursor populations in bone marrow. Fig. 6 directly compares the aeroallergen-induced bone marrow responses of offspring from treated versus untreated mothers. Firstly, while baseline output of pre-cDC and cDC was comparable between groups, there were small but significant increases in the resting GMP and MDP populations in the control offspring from treated mothers (Fig. 6C, D). However, the major treatment-associated differences were revealed by aeroallergen challenge, notably a consistent attenuation of the expansion in all precursor compartments spanning the MP–MDP stages which was observed in the challenged offspring from untreated mothers (Fig. 6A-D). Secondly, at the end of this developmental spectrum, the post challenge depletion of bone marrow cDC reserves that occurs in the offspring of untreated mothers was significantly attenuated (Fig. 6E, F), consistent with the reduced draw on this pool resulting from reduced recruitment to airway mucosa and ADLN in the OM85-treated group (Fig. 5A, B).

**Fig. 6.**
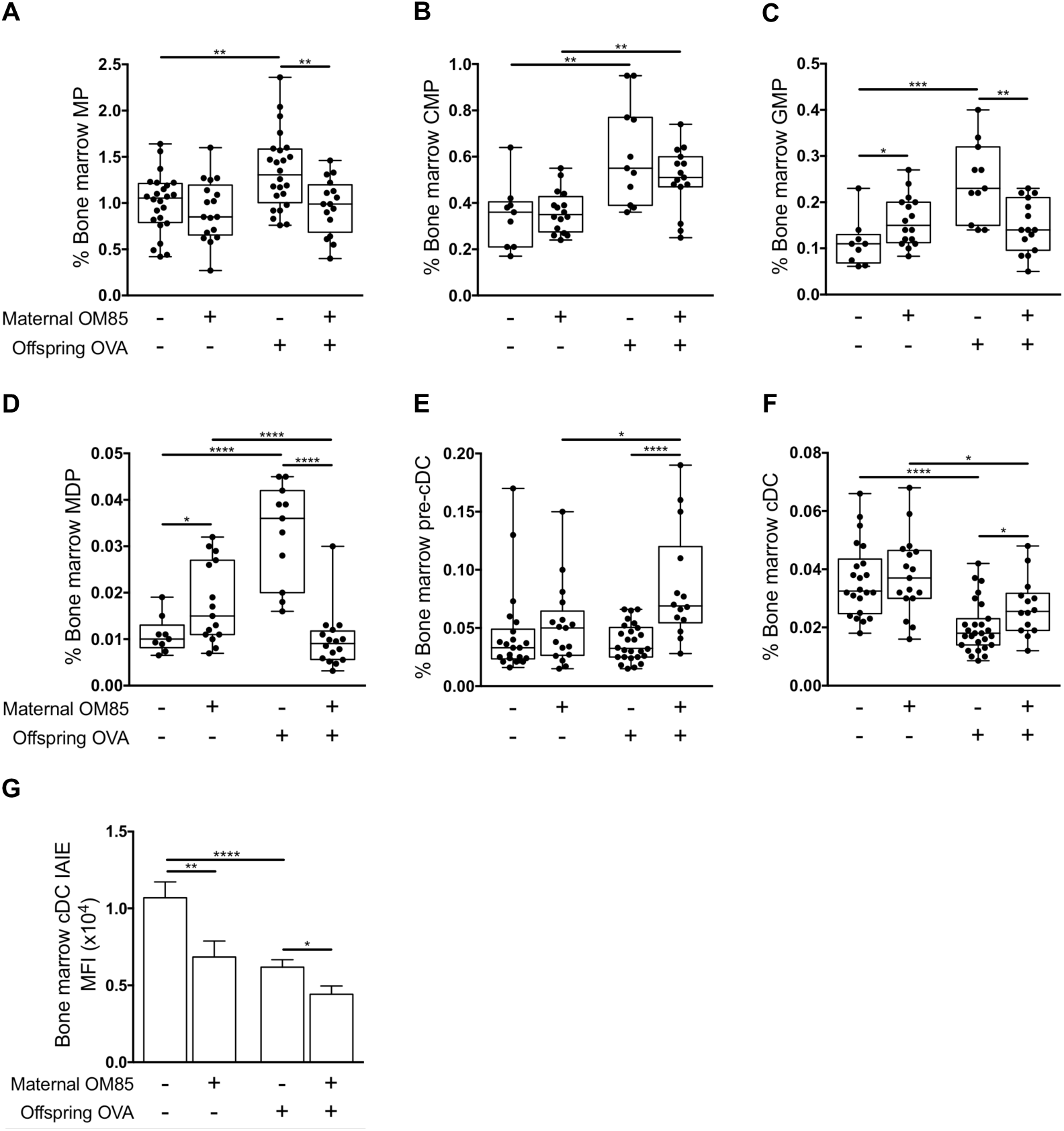
Attenuated bone marrow response following maternal OM85 pre-treatment. Bone marrow populations of MPs (A), CMPs (B), GMPs (C), MDPs (D), pre-cDCs (E), cDCs (**F**) and MFI of IAIE expression on cDCs (**G**) in OVA sensitised and aerosol challenged offspring versus naïve controls from OM85 treated/untreated mothers. Data displayed as box and whisker plot showing min to max or bar graph showing mean ± SEM of n=14 independent experiments. Data points represent individual animals. Statistical significance was determined using Student’s t-test or Mann Whitney test and presented as *p<0.05, **p<0.01, ***p<0.001, ****p<0.0001.

It is additionally pertinent to note that expression levels of IAIE on cDC were reduced in the treated group both at baseline and in particular post-challenge (Fig. 6G), suggesting maintenance of a more tightly regulated/quiescent functional state, mirroring the picture seen for trachea and ADLN cDC in Fig. 5. A similar pattern was also observed in relation to the bone marrow pDC reservoir (data not shown).

### Maternal OM85 treatment effects at earlier ages

The data above pertains to animals sensitized at 3 weeks and challenged/sacrificed at 6 weeks. In the studies presented in Fig. 7, we assessed the extent to which treatment effects on DC populations were demonstrable at younger ages. Looking firstly at age 3 weeks, we observed that the offspring of OM85-treated mothers displayed higher numbers CD11b^+^ and CD103^+^ cDC in lung tissue at baseline (Fig. 7A, B) and higher levels of attendant IAIE expression (Fig. 7C), consistent with treatment-mediated acceleration of postnatal maturation of DC networks in the respiratory tract. In a preliminary experiment we also compared total cDC yields in GM-CSF-driven bone marrow cultures derived from the same animals, and the increased yields from the treated group (Fig. 7D) again point to the marrow as the likely primary site of action of maternal OM85 treatment.

**Fig. 7.**
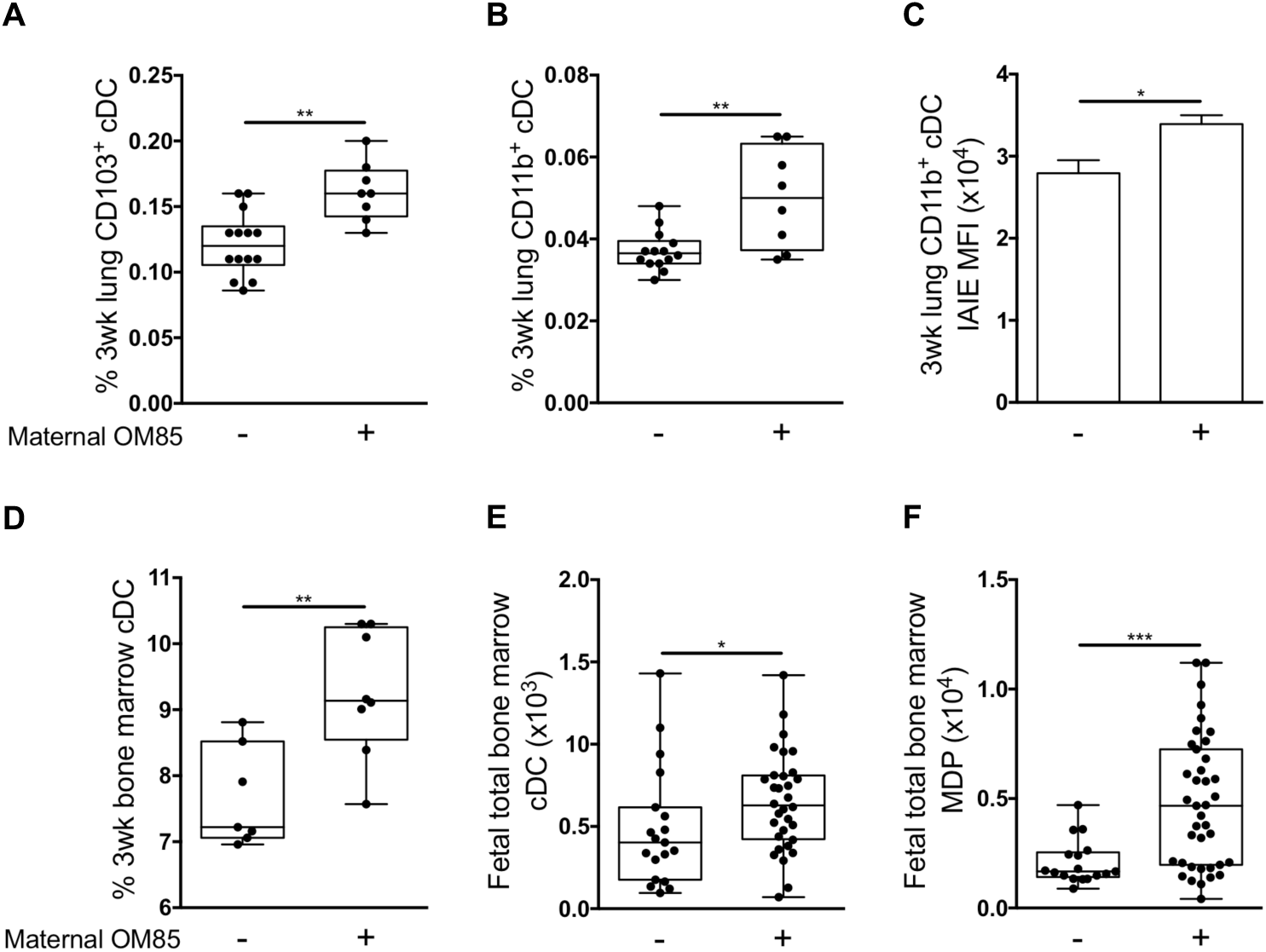
Maternal OM85 treatment boost early life development of the myeloid compartment. Peripheral lung CD103^+^ (**A**) and CD11b^+^ (**B**) cDC populations within 3-week old naïve offspring from OM85 treated/untreated mothers. (**C**) MFI of IAIE expression on peripheral lung CD11b^+^ cDCs from 3-week old naïve offspring from OM85 treated/untreated mothers. (**D**) Frequency of cDCs within *in vitro* 7-day bone marrow culture with GM-CSF from 3-week old naïve offspring from OM85 treated/untreated mothers. Absolute numbers of fetal bone marrow cDCs (**E**) and MDPs (**F**) collected at gestation day 18.5 from OM85 treated/untreated mothers. Data displayed as box and whisker plot showing min to max or bar graph showing mean ± SEM of n=5 independent experiments. Statistical significance was determined using Student’s t-test or Mann Whitney test and presented as *p<0.05, **p<0.01, ***p<0.001.

To further extend this finding, we characterised the progenitor pool within freshly harvested fetal bone marrow at 18.5 days gestation, 24 hours following the last maternal OM85 oral dose. The marked increase in total bone marrow cDC (Fig. 7E) accompanied by parallel expansion in the upstream MDP compartment in the treated group (Fig. 7F) is consistent with the conclusion that the bone marrow is a direct target for OM85 treatment effects.

### OM85-mediated attenuation of the responsiveness of bone marrow DC precursors to environmental inflammatory stimuli: validation of OM85 treatment effects in an independent inflammatory model

In the experiments illustrated in Fig. 8 we cultured bone marrow from 6-week old offspring of OM85-treated/untreated mothers in GM-CSF-enriched medium for 7 days, adding the archetypal pro-inflammatory agent bacterial LPS to half the cultures for the last 24 hours. Comparison of resultant activation levels of cDC by surface expression of IAIE (Fig. 8A) and the costimulator CD86 (Fig. 8B) indicated marked attenuation of upregulation of these function-associated markers, consistent with enhanced capacity for homeostatic regulation of inflammatory responses in general in cDC from the treated group.

**Fig. 8.**
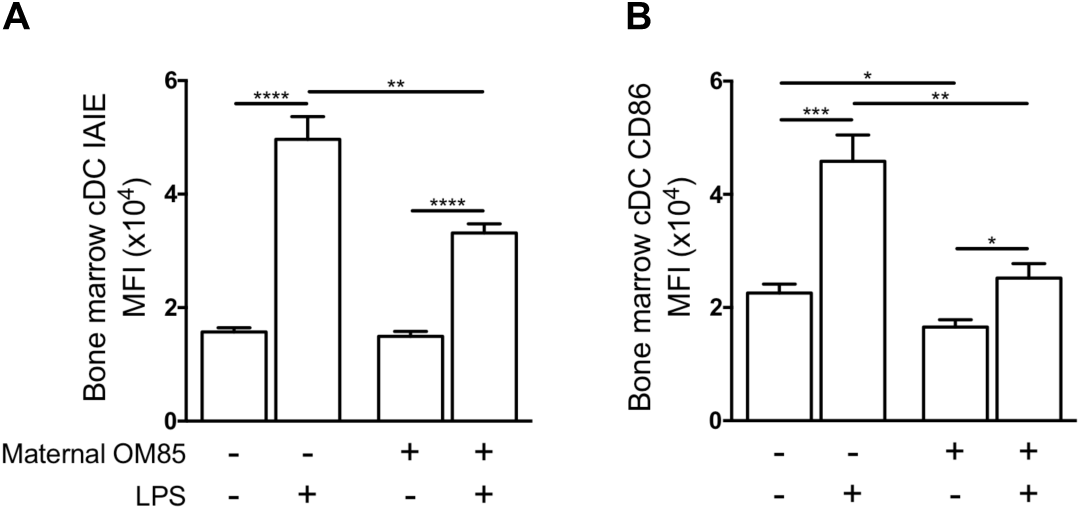
Validation of maternal OM85 protection via *in vitro* LPS stimulation. MFI of IAIE (**A**) and CD86 (**B**) expression on bone marrow cDCs cultured for 7 days in the presence of GM-CSF ± 10μg lipopolysaccharide for 24 hours. Data presented for naïve offspring from OM85 treated/untreated mothers and displayed as bar graph showing mean ± SEM of n=14 independent experiments. Statistical significance was determined using Student’s t-test or Mann Whitney test for intergroup comparisons and Paired Student’s t-test or Wilcoxon signed-rank test for intragroup comparisons and presented as *p<0.05, **p<0.01, ***p<0.001, ****p<0.0001.

## DISCUSSION

In this experimental asthma model, allergen challenge of early-life sensitized animals via aerosol triggers accumulation of a Th2-associated inflammatory cell infiltrate in respiratory tract tissues, and ensuing AHR. Consistent with earlier reports^15, 16^, prominent within these infiltrates were activated CD4^+^ T^m^effs and Treg cell populations, with the accompanying buildup of an expanded population of cDC and their transition (especially in the mucosa) from the passive/antigen surveillance phenotype (IAIE^low^) to a functionally mature (IAIE^high^) state. We also demonstrate for the first time that these rapid changes in the population dynamics of cDC in the challenged airways of sensitized animals are accompanied by concomitant depletion of committed cDC from bone marrow, and parallel (presumably compensatory) expansion of upstream multipotent precursor compartments.

We further demonstrate that susceptibility to this aeroallergen-induced asthma-like response in the airways is markedly attenuated in young animals born to mothers given repeated doses of OM85 during pregnancy. Accompanying this acquired resistant state is increased capacity for expansion and functional activation of Tregs in the airway mucosa in response to aeroallergen challenge, with parallel attenuation of local cDC recruitment, activation and trafficking to ADLN. Moreover, underpinning these treatment-related effects in respiratory tract associated tissues are a series of parallel changes in bone marrow DC progenitor populations at various stages of DC commitment, which are collectively consistent with the reduced draw on bone marrow cDC reserves in the treated group as a result of more effective control of the inflammatory milieu within the challenged mucosa. The key question is whether these changes in bone marrow of offspring from treated mothers are primary or secondary in this process.

In this regard, we^45^ and others^44^ have previously demonstrated that oral OM85 treatment of rodents can directly promote generation in gut-associated tissues of mucosal-homing DC that can bolster systemic natural Treg populations (including those in the airway mucosa) at baseline. However in the current model, airway mucosal Treg density and functional phenotype do not differ between OM85- and non-treated groups at baseline, and the stimulatory effects of treatment are only evident following aeroallergen challenge (Fig. 4C-K). We posit therefore that OM85 treatment has affected the functionality of the airway mucosal cDC that are responsible for programming Treg:Teff balance in the aerosol induced T-cell response, prior to their migration into the airway mucosa i.e. at bone marrow precursor stage. Several lines of indirect evidence support the possibility of direct OM85 treatment effects in bone marrow: (i) baseline IAIE expression on 6-week old bone marrow cDC is significantly reduced in offspring from treated mothers (Fig. 6G), and corresponding GMP and MDP precursor compartments are expanded in the same animals (Fig. 6C, D); (ii) at age 3 weeks when airway mucosal cDC networks are normally developmentally compromised with respect to baseline density, offspring from treated mothers display higher frequency of CD103^+^ and CD11b^+^ cDC in lung tissue digests (Fig. 7A, B) and higher cDC yields from bone marrow cultures (Fig. 7D); (iii) the frequency of cDC and their MDP precursors in fetal bone marrow are already increased in those from treated mothers by late gestation (Fig. 7E, F). However, the strongest evidence for treatment related effects at the DC precursor stage comes from the studies on cDC from 6-week old GM-CSF-driven 7-day bone marrow cultures (Fig. 8). The introduction of the archetypal pro-inflammatory stimulus bacterial LPS for the final 24 hours of these cultures triggers ultra-high expression on cDC from untreated controls of both IAIE and CD86, at levels likely to result in potentially pathologic T-cell hyperstimulation. However, this upregulation was more tightly controlled in cDC from offspring of treated mothers, suggesting enhanced capacity to maintain homeostasis and avoid bystander damage to host tissues during immunoinflammatory responses to environmental stimuli.

We acknowledge several limitations to this study. Firstly, we have not addressed the question of whether OM85 treatment influences susceptibility to primary allergic sensitization to inhalant allergens, and this merits future investigation, given that this process has also been shown to be controlled by ADLN-derived Tregs^46^. Secondly, we have no information on the mechanism of transmission of orally delivered OM85-associated signals to the bone marrow. Earlier studies from our group^45^ and others^44^ have demonstrated local activation of both T-cells and myeloid cells in the gut wall and associated lymphoid tissues following OM85 feeding, and it is possible that trafficking of representatives of either or both of these populations, or transmission of soluble signals generated at these sites, may be involved. Likewise, information on the gene target(s) in fetal bone marrow DC progenitors is not available, and will be the subject of future studies, along with possible effects on fetal thymus. Additionally, OM85 dosing of mothers in this study spanned only the second half of gestation, and future studies need to investigate the impact of extending feeding to earlier stages.

Our goal in this study was to provide a scientific rationale for subsequent use of OM85 during pregnancy in human mothers whose progeny are at risk of postnatal development of persistent atopy and/or asthma. The prime initial candidates for therapy in this regard are atopic asthmatic mothers^47, 48^. The severity of asthma symptomatology in this group is exaggerated during pregnancy^49-51^, likely as a consequence of the generalized Th2-skewing of immune functions associated with the pregnant state, and asthma exacerbations during pregnancy further increase asthma risk for their offspring^52^. Moreover, susceptibility to respiratory infections exemplified by influenza virus and its associated symptom severity is likewise increased during pregnancy^53, 54^, and the latter (along with bacterial infections) is a risk factor for fetal growth restriction^55^ which in turn is associated with increased risk for postnatal development of a range of non-communicable diseases including asthma^56, 57^. In this regard, our forerunner studies on OM85 use in pregnant mice have demonstrated strong protection against the effects of pathogen-associated challenge during pregnancy employing high dose LPS or live influenza, both with respect to preservation of pregnancy *per se,* and maintenance of normal maternal weight gain and associated fetal growth trajectories^13^. In this system the principal treatment effect of OM85 involved selective attenuation within gestational tissues of the intensity of the pro-inflammatory (particularly TNFα/IL-1/IL-6) components of the myeloid innate immune response to pathogen, with parallel preservation of vigorous Type 1 IFN-associated host defense networks^13^. Moreover, we^45^ and others^44^ have also previously demonstrated marked attenuation of airways inflammation in OM85-treated adult animals in models of experimental atopic asthma. On this basis, it can be argued that OM85 use during pregnancy has potential direct short-term benefits to mothers, as well as long-term benefits to their offspring in regards to reduced susceptibility to asthma development. The latter may also include enhanced resistance to early postnatal respiratory infections, given the OM85-associated effects demonstrated above on increased airway mucosal cDC numbers and IAIE expression at baseline in 3-week old weanlings, and studies are in progress to test this possibility.

A series of additional steps are required before progression to human trials with OM85 in pregnancy. The first involves an independent assessment by regulators of relevant safety issues. In this regard there is a wide body of data available on safe use during pregnancy in experimental animals, including our own^13^. Direct human safety data for pregnancy is not presently available. However, there is a >30-year history of safe use in non-pregnant adult humans and children down to age 6 months, and this has proven sufficient for relevant US^58^ (NIH) and Australian^59^ (NHMRC) authorities to endorse and publicly fund multicenter clinical trials in infants, targeting protection against wheezing symptoms (including those associated with early infections) and subsequent asthma development. In this regard, it is evident that airways inflammation associated with infections and inhalant allergy act in synergy to drive asthma pathogenesis during childhood^32^, and moreover that the earlier these episodic inflammatory events commence after birth, the greater is the risk for subsequent asthma^60-62^. This provides a compelling argument for development of protective therapeutic strategies that can reduce susceptibility to either or both of these environmental stressors, from birth onwards, and the possibility that this may be achievable prenatally with a readily available therapeutic such as OM85 merits further detailed investigation.

## MATERIALS AND METHODS

### Study Design

The objective of this pre-clinical BALB/c study was to identify a safe therapeutic immune modulating agent that could replicate the microbial-induced protection of offspring from allergic asthma observed on European farms and elucidate the underlying mechanism(s)-of-action promoting protection. We hypothesised that maternal OM85 treatment during gestation would alter the responsiveness and functional maturation of T cell and dendritic cell populations within the airways of antigen-sensitised and aerosol challenged offspring, thereby reducing offspring susceptibility to experimental asthma. However, observations made in the bone marrow revealed that the airways response may be secondary to upstream attenuation of the bone marrow compartment following maternal OM85 treatment. Differential cell counts, passive cutaneous anaphylaxis assays, airways hyperresponsiveness, flow cytometry and *in vitro* cell culture was used in combination to characterise the systemic response within offspring. Passive cutaneous anaphylaxis *in vivo* assays were performed using Sprague Dawley rats and all other *in vivo* and *in vitro* experiments were performed using BALB/c mice. Pregnant mice were selected at random for OM85 (400mg.kg^-1^) treatment during gestation. Offspring samples size was determined based on maternal n≥6. Experiments were independently replicated to a minimum of n=4 as outlined in corresponding figure legends. BAL differential cell counts were performed blind. All data was included unless technical issues during flow cytometry sample acquisition were deemed to have compromised the sample integrity.

### Animals

Specified pathogen-free BALB/c mice and Sprague Dawley (SD) rats were obtained from the Animal Resource Centre (Murdoch, WA, Australia). All animals were housed under specified pathogen-free conditions at the Telethon Kids Institute Bioresources Facility, with a 12h:12h light-dark cycle and access to an ovalbumin-free diet and water *ad libitum.* All animal experiments were approved and performed in accordance with the Telethon Kids Institute Animal Ethics Committee and the NHMRC guidelines for use of animals for scientific research. In-house bred BALB/c offspring of both sexes were used in these studies.

### Time-mated pregnancies

Female BALB/c mice 8 – 12 weeks of age were time-mated with male studs between 8 – 26 weeks of age. Male studs were housed separately in individual cages. One-to-two females were housed in an individual male cage overnight, with the presence of a vaginal plug the following morning used as an indicator of mating. The day of vaginal plug detection was designated gestation day (GD) 0.5.

### Maternal OM85 treatment protocol

Based on previously optimised dosing concentrations^13, 45^, time-mated pregnant female BALB/c mice selected at random received daily feeding of lyophilised OM85 (OM Pharma, Geneva, Switzerland) reconstituted in phosphate-buffered saline (PBS) to a concentration of 400mg.kg^-1^ body weight for the second half of gestation (GD9.5-17.5). Controls were left untreated. All treatments were performed from a single batch of OM85 (Batch no. 1812162).

### Offspring antigen sensitisation and aerosol challenge

Offspring from OM85 treated or naïve mothers were experimentally sensitised to ovalbumin (OVA) at the age of 21 and 35 days via intraperitoneal (i.p.) inoculation of 20μg OVA (grade V; Sigma-Aldrich, MO, USA) emulsified in 1.3mg aluminium hydroxide (Alu-Gel-S, SERVA, Heidelberg, Germany) in a total volume of 200μl. On days 42, 43 and 44, sensitised offspring were exposed OVA aerosol challenge (1% weight/volume in PBS) for 30 minutes, delivered via ultrasonic nebuliser. Mice for airways hyperresponsiveness assessment received a single OVA aerosol challenge on day 42 for 30 minutes. All experimental mice were sacrificed 24 hours post final aerosol.

### Measurement of airway hyperresponsiveness

Airway hyperresponsiveness (AHR) to inhaled methacholine (MCh) was assessed 24 hours after a single OVA (1% weight/volume in PBS) aerosol on day 42 following pre-sensitisation. The low-frequency forced oscillation technique (LFOT) was used to measure respiratory system input impedance (Z_rs_), as determined by previously optimised protocols^63^. Briefly, BALB/c mice were anesthetised (40% ketamine 100mg.ml^-1^, 10% xylazine 20mg.ml^-1^, 50% saline; 1% body weight), tracheotomised and ventilated (Legacy flexiVent, SCIREQ, Montreal, Canada) at 450 breaths/min with a tidal volume of 8ml.kg^-1^ and 2cmH_2_0 positive end-expiratory pressure (PEEP). Lung volume history was standardised for each individual mouse prior to measurement of experimental lung mechanics. Z_rs_ was measured during 16-second periods of apnea using a signal containing 19 mutually prime sinusoidal frequencies ranging from 0.25 to 19.625 Hz. The constant-phase model was fit to Z_rs_ in order to calculate changes in airways resistance (R_aw_). Ventilated BALB/c mice had 5x baseline measurements recorded, with 10-second aerosol challenge of saline followed by semi-log-fold increasing dose concentrations of MCh ranging from 0.1 – 30mg.ml^-1^ to assess for AHR. Five LFOT measurements were recorded after each MCh dose at 1 minute intervals. Dose-response curves were generated using the maximum response recorded for R_aw_.

### Tissue collection

***Fetal.*** Pregnant BALB/c mice were sacrificed on GD18.5. Both horns of the uterus were removed and fetuses were sacrificed via decapitation. Hind legs were removed, cleaned of remaining tissue and stored in cold PBS + 0.1% BSA. Dead fetuses were excluded. ***3-week old offspring.*** Offspring were sacrificed as 21 days of age. Lungs were perfused via cardiac flush with 2ml cold PBS + 0.1% BSA. Peripheral lung and the femur and tibia from both hind legs were collected. ***6-week old offspring.*** Offspring were sacrificed at 45 days of age. Lungs were perfused via cardia flush with 2ml cold PBS + 0.1% BSA. Parathymic and mediastinal (airways draining) lymph nodes (ADLN), trachea, peripheral lung and the femur and tibia from both hind legs were collected. Blood was collected via cardiac puncture at time of autopsy.

### Passive cutaneous anaphylaxis IgE assay

*In vivo* passive cutaneous anaphylaxis assays were performed using male Sprague Dawley rats >10 weeks of age. Individual BALB/c serum samples were prepared as serial 1:2 dilutions with a final sample volume of 55μl SD rats were anaesthetised via i.p. injection of 4ml 5.71% chloral hydrate (Sigma-Aldrich, MO, USA) in PBS. Once anesthetised, rats had their back closely shaved to remove all hair and 50μl of each sample was subcutaneously injected down the back using a 1ml 29g insulin syringe. 24 hours later, rats were anaesthetised with chloral hydrate and intravenously (i.v.) injected with 2ml of a 1:1 antigen-dye solution containing 4mg.ml^-1^ OVA in PBS and 1% Evans blue dye (Sigma-Aldrich, MO, USA). Blue subcutaneous injection sites after 15-30 minutes indicate serum samples positive for OVA-specific IgE. The highest positive serum dilution for each sample was recorded and animals were euthanised with 600μl Lethabarb i.v.

### Single cell suspension preparation

#### Airways tissue and fetal bone marrow

ADLN, trachea, peripheral lung and fetal bone marrow single cell suspensions were prepared by mincing excised tissue/bone into smaller pieces and resuspending in 10ml GKN (11mM D-glucose, 5.5mM KCl, 137 mM NaCl, 25mM Na_2_HPO_4_) + 10% fetal calf serum (FCS; Serana, Bunbury, WA, Australia) with collagenase IV (Worthington Biochemical Corporation, Lakewood, NJ, USA) and DNase (Sigma-Aldrich, MO, USA) for enzymatic digestion at 37°C under gentle agitation for 30 minutes (ADLN and trachea), 60 minutes (fetal bone marrow) or 90 minutes (peripheral lung). Following digestion, tissues were disaggregated via manual pipetting and filtered through sterile cotton wool columns. Cell suspensions were centrifuged and pellet resuspended in red blood cell (RBC; 17mM Tris-HCl, 0.14M NH_4_Cl at pH 7.2) lysis buffer for 3 minutes. Cell were washed with cold PBS and pelleted. Supernatant was removed and pellet resuspended in PBS + 0.1% BSA for total cell counts. ***3- and 6-week old bone marrow***. Long bones were flushed with 10ml GKN + 5% FCS using a 25g needle. Cells were disaggregated by manual pipetting and filtered through a sterile cotton wool column. Filtered cells were washed with GKN + 5% FCS and centrifuged at 1800rpm for 8 minutes at 4°C. Supernatant was removed and pellet resuspended in RBC lysis buffer for 5 minutes. Cells were washed in cold PBS, centrifuged and pellet resuspended in PBS + 0.1% BSA for total cell counts.

### Bronchoalveolar lavage and differential cell counts

Bronchoalveolar lavage (BAL) fluid was collected via tracheal cannula flushing the lungs three times with 800μl cold PBS + 0.1% BSA. BAL cells were resuspended in 300μl RBC lysis buffer for 4 minutes. Cells were washed with cold PBS, spun and pellets resuspended in 100μl cold PBS + 0.1% BSA for counting. BAL samples were counted with Trypan Blue (LabChem; Thermo Fisher Scientific, MA, USA) using a haemocytometer and counting to a minimum of 100 leukocytes. 1x10^5^ cells for each individual sample were spun onto Superfrost^®^ Plus microscope slides (LabServ; Thermo Fisher Scientific, MA, USA). Cytospin cell preparations were stained using Diff-Quik (Rapid Stain Kit; Perth Scientific, WA, Australia) and differential cell counts performed by counting ≥300 cells per cytospin.

### Bone marrow cultures

#### 3-week offspring

Single-cell bone marrow suspensions (previously described) were washed with 20ml cold PBS + 0.1% BSA and centrifuged at 1800rpm for 8 minutes at 4°C. Cells were resuspended in RPMI-10 Complete media (RPMI 1640, 10% FCS, 2mM L-glutamine, 50μM 2-β-mercaptoethanol (Sigma-Aldrich, MO, USA), 5μg.ml^-1^ gentamycin (Pfizer, NY, USA) and 10ng.ml^-1^ GM-CSF) at a concentration of 8x10^5^ cells.ml^-1^. 1ml aliquots were seeded onto 24-well treated cell culture plates and incubated at 37°C and 5% CO2 in a water jacketed incubator. At 48 hours, culture media was aspirated and wells were washed with 1ml RPMI supplemented with 2mM L-glutamine, 50μM 2-β-mercaptoethanol and 5μg.ml^-1^ gentamycin. Wash was aspirated and 1ml of fresh RPMI-10 Complete media was added to wells. After 6 days, cells were harvested and wells washed twice with 500μl RPMI-10. Cells were centrifuged and pellets resuspended in 500μl RPMI-10 to perform total cell counts. Following counts, cells were resuspended at a density of 1x10^6^ cells.ml^-1^ in RPMI-10 Complete media and 1ml aliquots were re-seeded on a 24-well treated cell culture plate. After 24 hours, cells were harvested for flow cytometric phenotypic analysis. ***6-week offspring***. Culture days 1-5 were as described for 3-week offspring. On day 6, cells were harvested into 15ml conical tubes and wells washed twice with 500μl RPMI-10. Cells were centrifuged and pellets resuspended in 500μl RPMI-10 to perform total cell counts. Following counts, cells were resuspended at a density of 1x10^6^ cells.ml^-1^ in RPMI-10 Complete media. 1ml aliquots were re-seeded on a 24-well treated cell culture plate and 1ng.ml^-1^ lipopolysaccharide (LPS) was added to each well. Cells were cultured in the presence of LPS for 24 hours. After 24 hours, cells were harvested for flow cytometric phenotypic analysis.

### Flow cytometry

Single-cell suspensions (as per above) were used for all immunostaining. Panels of monoclonal antibodies were developed to enable identification of airways myeloid (IA/IE-AF647, F4/80-FITC, CD11c-AF700, CD11b-V500, Ly6G/C-APC Cy7, B220-PE CF594, CD8α-Bς650, CD40-BV421, CD45-PerCP, CD103-PE, CD86-PE Cy5 and NKp46-PE Cy7), airways T cell (CD3-FITC, CD4-V500, CD8α-BV650, CD25-APC Cy7, CD45-PerCP, CD69-PE Cy7, CTLA-4-BV421, Ki67-AF700 and FoxP3-PE), bone marrow myeloid (CD3-FITC, CD11b-BV510, CD11c-BV711, CD19-APC Cy7, Gr-1-BV605, B220-PerCP Cy5.5, NKp46-Pe Cy7, SIRP-α-APC, IA/IE-BV421 and F4/80-BV785) and bone marrow hematopoietic stem and progenitor cell (CD2-BV605, CD3-BV605, CD4-BV605, CD5-BV605, CD8α-BV605, CD19-BV605, B220-BV605, Gr-1-BV605, Ter119-BV605, CD16/32-PerCP Cy5.5, CD34-FITC IL-7Rα,-PE Cy7 Flt-3-PE, c-Kit-APC Cy7, Sca-1-BV521, CX_3_CR1-APC and NKG2D-BV711) populations. All reagents were supplied by BD Biosciences, San Jose, CA, USA or BioLegend, San Diego CA, USA unless otherwise stated. Intracellular staining for FoxP3, CTLA-4 and Ki67 was performed using a FoxP3 intracellular staining buffer set (eBiosciences, San Diego, CA, USA). Acquisition was performed on a four-laser LSR Fortessa™ (BD Bioscience, San Jose, CA, USA). All samples were kept as individuals and not pooled. Immune cell phenotyping was analysed using FlowJo^®^ software (Version 10.1, Tree Star, Sanford, CA, USA) and associated gating strategies are outline in Fig. S4-S6.

### viSNE analysis

ADLN, trachea and peripheral lung FCS files, with software compensation applied, were uploaded to the Cytobank platform (Cytobank Mountain View, CA, USA) and analysed using established methods^64, 65^. The software transformed the data to arcsinh scales. Equal cell numbers were used from each FCS file. Antibodies listed as per above were used for T cell subset identification to create viSNE maps using a total of 100,000 (ADLN and peripheral lung) or 3,000 (trachea) cells per sample.

### Statistical analysis

Statistical analysis and graphing was performed using GraphPad Prism (GraphPad software; version 7.0a). Statistical significance of p<0.05 was considered significant.

## REFERENCES

1 von Mutius E, Radon K, Living on a Farm: Impact on Asthma Induction and Clinical Course. Immunol Allergy Clin North Am 28, 631–647 (2008).

2 Ege MJ, Mayer M, Normand A-C, Genuneit J, Cookson WOCM, Braun-Fahrländer C, Heederik D, Piarroux R, von Mutius E, Exposure to Environmental Microorganisms and Childhood Asthma. N Engl J Med 364, 701–709 (2011).

3 Stein MM, Hrusch CL, Gozdz J, Igartua C, Pivniouk V, Murray SE, Ledford JG, Marques dos Santos M, Anderson RL, Metwali N, Neilson JW, Maier RM, Gilbert JA, Holbreich M, Thorne PS, Martinez FD, von Mutius E, Vercelli D, Ober C, Sperling AI, Innate Immunity and Asthma Risk in Amish and Hutterite Farm Children. N Engl J Med 375, 411–421 (2016).

4 Schaub B, Liu J, Höppler S, Schleich I, Huehn J, Olek S, Wieczorek G, Illi S, von Mutius E, Maternal farm exposure modulates neonatal immune mechanisms through regulatory T cells. J Allergy Clin Immunol 123, 774–782.e775 (2009).

5 Schuijs MJ, Willart MA, Vergote K, Gras D, Deswarte K, Ege MJ, Madeira FB, Beyaert R, van Loo G, Bracher F, von Mutius E, Chanez P, Lambrecht BN, Hammad H, Farm dust and endotoxin protect against allergy through A20 induction in lung epithelial cells. Science 349, 1106–1110 (2015).

6 Holt PG, Sly PD, Environmental Microbial Exposure and Protection against Asthma. N Engl J Med 373, 2576–2578 (2015).

7 Ege MJ, Bieli C, Frei R, van Strien RT, Riedler J, Üblagger E, Schram-Bijkerk D, Brunekreef B, van Hage M, Scheynius A, Pershagen G, Benz MR, Lauener R, von Mutius E, Braun-Fahrländer C, the PSt, Prenatal farm exposure is related to the expression of receptors of the innate immunity and to atopic sensitization in school-age children. J Allergy Clin Immunol 117, 817–823 (2006).

8 Schaad UB, Mutterlein R, Goffin H, Immunostimulation with OM-85 in children with recurrent infections of the upper respiratory tract: a double-blind, placebo-controlled multicenter study. Chest 122, 2042–2049 (2002).

9 Razi CH, Harmanci K, Abaci A, Özdemir O, Hizli Ş, Renda R, Keskin F, The immunostimulant OM-85 BV prevents wheezing attacks in preschool children. J Allergy Clin Immunol 126, 763–769 (2010).

10 Collet JP, Shapiro P, Ernst P, Renzi T, Ducruet T, Robinson A, Effects of an immunostimulating agent on acute exacerbations and hospitalizations in patients with chronic obstructive pulmonary disease. The PARI-IS Study Steering Committee and Research Group. Prevention of Acute Respiratory Infection by an Immunostimulant. Am JRespir Crit Care Med 156, 1719–1724 (1997).

11 Orcel B, Delclaux B, Baud M, Derenne JP, Oral immunization with bacterial extracts for protection against acute bronchitis in elderly institutionalized patients with chronic bronchitis. Eur Respir J 7, 446–452 (1994).

12 Solèr M, Mutterlein R, Cozma G, Double-blind study of OM-85 in patients with chronic bronchitis or mild chronic obstructive pulmonary disease. Respiration 74, 26-32 (2007).

13 Scott NM, Lauzon-Joset JF, Jones AC, Mincham KT, Troy NM, Leffler J, Serralha M, Prescott SL, Robertson SA, Pasquali C, Bosco A, Holt PG, Strickland DH, Protection against maternal infection-associated fetal growth restriction: proof-of-concept with a microbial-derived immunomodulator. Mucosal Immunol 10, 789–801 (2017).

14 Holt PG, Upham JW, Sly PD, Contemporaneous maturation of immunologic and respiratory functions during early childhood: Implications for development of asthma prevention strategies. J Allergy Clin Immunol 116, 16–24 (2005).

15 Huh JC, Strickland DH, Jahnsen FL, Turner DJ, Thomas JA, Napoli S, Tobagus I, Stumbles PA, Sly PD, Holt PG, Bidirectional Interactions between Antigen-bearing Respiratory Tract Dendritic Cells (DCs) and T Cells Precede the Late Phase Reaction in Experimental Asthma: DC Activation Occurs in the Airway Mucosa but Not in the Lung Parenchyma. JExp Med 198, 19–30 (2003).

16 Stumbles PA, Thomas JA, Pimm CL, Lee PT, Venaille TJ, Proksch S, Holt PG, Resting respiratory tract dendritic cells preferentially stimulate T helper cell type 2 (Th2) responses and require obligatory cytokine signals for induction of Th1 immunity. J Exp Med 188, 2019–2031 (1998).

17 Vermaelen KY, Carro-Muino I, Lambrecht BN, Pauwels RA, Specific migratory dendritic cells rapidly transport antigen from the airways to the thoracic lymph nodes. J Exp Med 193, 51–60 (2001).

18 Akdis M, Verhagen J, Taylor A, Karamloo F, Karagiannidis C, Crameri R, Thunberg S, Deniz G, Valenta R, Fiebig H, Kegel C, Disch R, Schmidt-Weber CB, Blaser K, Akdis CA, Immune responses in healthy and allergic individuals are characterized by a fine balance between allergen-specific T regulatory 1 and T helper 2 cells. J Exp Meed 199, 1567–1575 (2004).

19 Strickland DH, Stumbles PA, Zosky GR, Subrata LS, Thomas JA, Turner DJ, Sly PD, Holt PG, Reversal of airway hyperresponsiveness by induction of airway mucosal CD4(+)CD25(+) regulatory T cells. J Exp Med 203, 2649–2660 (2006).

20 Cederbom L, Hall H, Ivars F, CD4+CD25+ regulatory T cells down-regulate costimulatory molecules on antigen-presenting cells. Eur J Immunol 30, 1538–1543 (2000).

21 Lewkowich IP, Herman NS, Schleifer KW, Dance MP, Chen BL, Dienger KM, Sproles AA, Shah JS, Köhl J, Belkaid Y, Wills-Karp M, CD4(+)CD25(+) T cells protect against experimentally induced asthma and alter pulmonary dendritic cell phenotype and function. J Exp Med 202, 1549–1561 (2005).

22 De Heer HJ, Hammad H, Soullié T, Hijdra D, Vos N, Willart MAM, Hoogsteden HC, Lambrecht BN, Essential role of lung plasmacytoid dendritic cells in preventing asthmatic reactions to harmless inhaled antigen. J Exp Med 200, 89–98 (2004).

23 Gill MA, Palucka AK, Barton T, Ghaffar F, Jafri H, Banchereau J, Ramilo O, Mobilization of plasmacytoid and myeloid dendritic cells to mucosal sites in children with respiratory syncytial virus and other viral respiratory infections. J Infect Dis 191, 1105–1115 (2005).

24 Holt PG, Haining S, Nelson DJ, Sedgwick JD, Origin and steady-state turnover of class II MHC-bearing dendritic cells in the epithelium of the conducting airways. J Immunol 153, 256–261 (1994).

25 McWilliam AS, Nelson D, Thomas JA, Holt PG, Rapid dendritic cell recruitment is a hallmark of the acute inflammatory response at mucosal surfaces. J Exp Med 179, 1331–1336 (1994).

26 McWilliam AS, Napoli S, Marsh AM, Pemper FL, Nelson DJ, Pimm CL, Stumbles PA, Wells TNC, Holt PG, Dendritic Cells Are Recruited into the Airway Epithelium during the Inflammatory Response to a Broad Spectrum of Stimuli. J Exp Med 184, 2429–2432 (1996).

27 Jahnsen FL, Strickland DH, Thomas JA, Tobagus IT, Napoli S, Zosky GR, Turner DJ, Sly PD, Stumbles PA, Holt PG, Accelerated antigen sampling and transport by airway mucosal dendritic cells following inhalation of a bacterial stimulus. J Immunol 177, 5861–5867 (2006).

28 Tschernig T, Debertin AS, Paulsen F, Kleemann WJ, Pabst R, Dendritic cells in the mucosa of the human trachea are not regularly found in the first year of life. Thorax 56, 427–431 (2001).

29 Heier I, Malmstrom K, Sajantila A, Lohi J, Makela M, Jahnsen FL, Characterisation of bronchus-associated lymphoid tissue and antigen-presenting cells in central airway mucosa of children. Thorax 66, 151–156 (2011).

30 Nelson DJ, McMenamin C, McWilliam AS, Brenan M, Holt PG, Development of the airway intraepithelial dendritic cell network in the rat from class II major histocompatibility (Ia)-negative precursors: differential regulation of Ia expression at different levels of the respiratory tract. J Exp Med 179, 203–212 (1994).

31 Nelson DJ, Holt PG, Defective regional immunity in the respiratory tract of neonates is attributable to hyporesponsiveness of local dendritic cells to activation signals. J Immunol 155, 3517–3524 (1995).

32 Holt PG, Sly PD, Viral infections and atopy in asthma pathogenesis: new rationales for asthma prevention and treatment. >Nat Med 18, 726 (2012).

33 Conrad ML, Ferstl R, Teich R, Brand S, Blumer N, Yildirim AO, Patrascan CC, Hanuszkiewicz A, Akira S, Wagner H, Holst O, von Mutius E, Pfefferle PI, Kirschning CJ, Garn H, Renz H, Maternal TLR signaling is required for prenatal asthma protection by the nonpathogenic microbe Acinetobacter lwoffii F78. J Exp Med 206, 2869–2877 (2009).

34 Strickland DH, Thomas JA, Mok D, Blank F, McKenna KL, Larcombe AN, Sly PD, Holt PG, Defective aeroallergen surveillance by airway mucosal dendritic cells as a determinant of risk for persistent airways hyper-responsiveness in experimental asthma. Mucosal Immunol 5, 332–341 (2012).

35 Plantinga M, Guilliams M, Vanheerswynghels M, Deswarte K, Branco-Madeira F, Toussaint W, Vanhoutte L, Neyt K, Killeen N, Malissen B, Hammad H, Lambrecht Bart N, Conventional and Monocyte-Derived CD11b(+) Dendritic Cells Initiate and Maintain T Helper 2 Cell-Mediated Immunity to House Dust Mite Allergen. Immunity 38, 322–335 (2013).

36 Kool M, Soullie T, van Nimwegen M, Willart MA, Muskens F, Jung S, Hoogsteden HC, Hammad H, Lambrecht BN, Alum adjuvant boosts adaptive immunity by inducing uric acid and activating inflammatory dendritic cells. J Exp Med 205, 869-882 (2008).

37 Hammad H, Plantinga M, Deswarte K, Pouliot P, Willart MAM, Kool M, Muskens F, Lambrecht BN, Inflammatory dendritic cells—not basophils—are necessary and sufficient for induction of Th2 immunity to inhaled house dust mite allergen. J Exp Med 207, 2097–2111 (2010).

38 Denburg JA, Sehmi R, Saito H, Pil-Seob J, Inman MD, O’Byrne PM, Systemic aspects of allergic disease: Bone marrow responses. J Allergy Clin Immunol 106, S242–S246 (2000).

39 Akashi K, Traver D, Miyamoto T, Weissman IL, A clonogenic common myeloid progenitor that gives rise to all myeloid lineages. Nature 404, 193–197 (2000).

40 Iwasaki H, Mizuno S-i, Mayfield R, Shigematsu H, Arinobu Y, Seed B, Gurish MF, Takatsu K, Akashi K, Identification of eosinophil lineage–committed progenitors in the murine bone marrow. J Exp Med 201, 1891–1897 (2005).

41 Fogg DK, Sibon C, Miled C, Jung S, Aucouturier P, Littman DR, Cumano A, Geissmann F, A clonogenic bone marrow progenitor specific for macrophages and dendritic cells. Science 311, 83–87 (2006).

42 Auffray C, Fogg DK, Narni-Mancinelli E, Senechal B, Trouillet C, Saederup N, Leemput J, Bigot K, Campisi L, Abitbol M, Molina T, Charo I, Hume DA, Cumano A, Lauvau G, Geissmann F, CX(3)CR1(+) CD115(+) CD135(+) common macrophage/DC precursors and the role of CX(3)CR1 in their response to inflammation. J Exp Med 206, 595–606 (2009).

43 del Rio ML, Rodriguez-Barbosa JI, Bolter J, Ballmaier M, Dittrich-Breiholz O, Kracht M, Jung S, Forster R, CX3CR1+ c-kit+ bone marrow cells give rise to CD103+ and CD103-dendritic cells with distinct functional properties. J Immunol 181, 6178–6188 (2008).

44 Navarro S, Cossalter G, Chiavaroli C, Kanda A, Fleury S, Lazzari A, Cazareth J, Sparwasser T, Dombrowicz D, Glaichenhaus N, Julia V, The oral administration of bacterial extracts prevents asthma via the recruitment of regulatory T cells to the airways. Mucosal Immunol 4, 53–65 (2011).

45 Strickland DH, Judd S, Thomas JA, Larcombe AN, Sly PD, Holt PG, Boosting airway T-regulatory cells by gastrointestinal stimulation as a strategy for asthma control. Mucosal Immunol 4, 43–52 (2011).

46 Holt PG, Strickland DH, Wikstrom ME, Jahnsen FL, Regulation of immunological homeostasis in the respiratory tract. Nat Rev Immunol 8, 142–152 (2008).

47 Tariq SM, Matthews SM, Hakim EA, Stevens M, Arshad SH, Hide DW, The prevalence of and risk factors for atopy in early childhood: A whole population birth cohort study. J Allergy Clin Immunol 101, 587–593 (1998).

48 Litonjua AA, Carey VJ, Burge HA, Weiss ST, Gold DR, Parental history and the risk for childhood asthma. Does mother confer more risk than father? Am J Respir Crit Care Med 158, 176–181 (1998).

49 Schatz M, Harden K, Forsythe A, Chilingar L, Hoffman C, Sperling W, Zeiger RS, The course of asthma during pregnancy, post partum, and with successive pregnancies: a prospective analysis. J Allergy Clin Immunol 81, 509–517 (1988).

50 Schatz M, Dombrowski MP, Wise R, Thom EA, Landon M, Mabie W, Newman RB, Hauth JC, Lindheimer M, Caritis SN, Leveno KJ, Meis P, Miodovnik M, Wapner RJ, Paul RH, Varner MW, O’Sullivan MJ, Thurnau GR, Conway D, McNellis D, Asthma morbidity during pregnancy can be predicted by severity classification. J Allergy Clin Immunol 112, 283–288 (2003).

51 Murphy VE, Gibson P, Talbot PI, Clifton VL, Severe asthma exacerbations during pregnancy. Obstet Gynecol 106, 1046–1054 (2005).

52 Martel MJ, Rey É, Beauchesne MF, Malo JL, Perreault S, Forget A, Blais L, Control and severity of asthma during pregnancy are associated with asthma incidence in offspring: Two-stage case-control study. Eur Respir J 34, 579–587 (2009).

53 Dodds L, McNeil SA, Fell DB, Allen VM, Coombs A, Scott J, MacDonald N, Impact of influenza exposure on rates of hospital admissions and physician visits because of respiratory illness among pregnant women. CMAJ: Canadian Medical Association Journal 176, 463–468 (2007).

54 Jamieson DJ, Honein MA, Rasmussen SA, Williams JL, Swerdlow DL, Biggerstaff MS, Lindstrom S, Louie JK, Christ CM, Bohm SR, Fonseca VP, Ritger KA, Kuhles DJ, Eggers P, Bruce H, Davidson HA, Lutterloh E, Harris ML, Burke C, Cocoros N, Finelli L, MacFarlane KF, Shu B, Olsen SJ, H1N1 2009 influenza virus infection during pregnancy in the USA. The Lancet 374, 451–458 (2009).

55 McNeil SA, Dodds LA, Fell DB, Allen VM, Halperin BA, Steinhoff MC, MacDonald NE, Effect of respiratory hospitalization during pregnancy on infant outcomes. Am J Obstet Gynecol 204, S54–S57 (2011).

56 Barker DJ, Adult consequences of fetal growth restriction. Clin Obstet Gynecol 49, 270–283 (2006).

57 Xu X-F, Li Y-J, Sheng Y-J, Liu J-L, Tang L-F, Chen Z-M, Effect of low birth weight on childhood asthma: a meta-analysis. BMC Pediatr 14, 275 (2014).

58 Oral Bacterial Extract for the Prevention of Wheezing Lower Respiratory Tract Illness - ClinicalTrials.gov 2017. Available from: https://clinicaltrials.gov/ct2/show/NCT02148796.

59 ANZCTR - Registration 2008. Available from: https://www.anzctr.org.au/Trial/Registration/TrialReview.aspx?ACTRN=12608000475347.

60 Bisgaard H, Hermansen MN, Buchvald F, Loland L, Halkjaer LB, Bonnelykke K, Brasholt M, Heltberg A, Vissing NH, Thorsen SV, Stage M, Pipper CB, Childhood asthma after bacterial colonization of the airway in neonates. N Engl J Med 357, 1487–1495 (2007).

61 Teo SM, Mok D, Pham K, Kusel M, Serralha M, Troy N, Holt BJ, Hales BJ, Walker ML, Hollams E, Bochkov YA, Grindle K, Johnston SL, Gern JE, Sly PD, Holt PG, Holt KE, Inouye M, The infant nasopharyngeal microbiome impacts severity of lower respiratory infection and risk of asthma development. Cell host & microbe 17, 704-715 (2015).

62 Kusel MMH, de Klerk NH, Kebadze T, Vohma V, Holt PG, Johnston SL, Sly PD, Early-life respiratory viral infections, atopic sensitization, and risk of subsequent development of persistent asthma. J Allergy Clin Immunol 119, 1105–1110 (2007).

63 Zosky GR, Larcombe AN, White OJ, Burchell JT, Janosi TZ, Hantos Z, Holt PG, Sly PD, Turner DJ, Ovalbumin-sensitized mice are good models for airway hyperresponsiveness but not acute physiological responses to allergen inhalation. Clin Exp Allergy 38, 829–838 (2008).

64 Kotecha N, Krutzik PO, Irish JM, Web-based analysis and publication of flow cytometry experiments. Current protocols in cytometry Chapter 10, Unit10.17 (2010).

65 Diggins KE, Ferrell PB, Jr., Irish JM, Methods for discovery and characterization of cell subsets in high dimensional mass cytometry data. Methods 82, 55–63 (2015).

